# Development of a Conditional Plasmid for Gene Deletion in Non-Model *Fusobacterium nucleatum* strains

**DOI:** 10.1101/2024.09.09.612158

**Authors:** Peng Zhou, G C Bibek, Chenggang Wu

**Affiliations:** Department of Microbiology & Molecular Genetics, the University of Texas Health Science Center, Houston, TX, USA

**Keywords:** *Fusobacterium nucleatum*, *repA*, theophylline-responsible riboswitch, gene deletion

## Abstract

*Fusobacterium nucleatum* is an opportunistic pathogen with four subspecies: *nucleatum* (FNN), *vincentii* (FNV), *polymorphum* (FNP), and *animalis* (FNA), each with distinct disease potentials. Research on fusobacterial pathogenesis has mainly focused on the model strain ATCC 23726 from FNN. However, this narrow focus may overlook significant behaviors of other FNN strains and those from other subspecies, given the genetic and phenotypic diversity within *F. nucleatum*. While ATCC 23726 is highly transformable, most other *Fusobacterium* strains exhibit low transformation efficiency, complicating traditional gene deletion methods that rely on non-replicating plasmids. To address this, we developed a conditional plasmid system in which the RepA protein, essential for replication of a pCWU6-based shuttle plasmid, is controlled by an inducible system combining an *fdx* promoter with a theophylline-responsive riboswitch. This system allows plasmid replication in host cells upon induction and plasmid loss when the inducer is removed, forcing chromosomal integration via homologous recombination in the presence of the antibiotic thiamphenicol. We validated this approach by targeting the *galK* gene, successfully generating mutants in FNN (ATCC 23726, CTI-2), FNP (ATCC 10953), FNA (21_1A), and the closely related species *Fusobacterium periodonticum*. Incorporating a *sacB* counterselection marker in this conditional plasmid enabled the deletion of the *radD* gene in non-model strains. Interestingly, while *radD* deletion in 23726, 10953, and 21_1A abolished coaggregation with *Actinomyces oris*, the CTI-2 mutant retained this ability, suggesting the involvement of other unknown adhesins. This work significantly advances gene deletion in genetically recalcitrant *F. nucleatum* strains, enhancing our understanding of this pathogen.

**IMPORTANCE:** *Fusobacterium nucleatum* is implicated in various human diseases, including periodontal disease, preterm birth, and colorectal cancer, often linked to specific strains and reflecting the species’ genetic and phenotypic diversity. Despite this diversity, most genetic research has centered on the model strain ATCC 23726, potentially missing key aspects of other strains’ pathogenic potential. This study addresses a critical gap by developing a novel conditional plasmid system that enables gene deletion in genetically recalcitrant strains of *F. nucleatum*. We successfully deleted genes in the clinical strain CTI-2, the FNA strain 21_1A, and *F. periodonticum* for the first time. Our findings, particularly the varying behavior of the *radD* gene production in coaggregation across strains, underscore the complexity of *F. nucleatum* and the need for broader genetic studies. This work advances our understanding of *F. nucleatum* virulence at the strain level and provides a valuable tool for future bacterial genetics research.

## INTRODUCTION

*Fusobacterium nucleatum* is a Gram-negative, anaerobic bacterium predominantly found in the human oral cavity, particularly in the subgingival biofilm, where it is highly abundant (1, 2). This bacterium plays a crucial role in dental plaque formation by physically interacting with early and late colonizers, serving as a “bridge” through cell surface adhesins like RadD, Fap2, and FomA (3–6). Consequently, *F. nucleatum* is closely associated with the development of periodontitis. Beyond this, *F. nucleatum* is strongly linked to oral carcinogenesis (7) and, when disseminated beyond the oral cavity, to various systemic diseases like colorectal cancer, adverse pregnancy outcomes, inflammatory bowel disease, and rheumatoid arthritis (8, 9).

*F. nucleatum* exhibits considerable genetic and phenotypic diversity and is currently classified into four subspecies: *nucleatum* (FNN), *vincentii* (FNV), *animalis* (FNA), and *polymorphum* (FNP) (10). Comparative genomics reveals that each subspecies possesses unique gene clusters (11–13), which may explain why strains from different subspecies exhibit distinct phenotypes in biofilm formation and host interactions(14–16). These differences likely determine the roles of different subspecies in human health and disease. For instance, FNP and FNA are the most abundant subspecies within the oral cavity, with FNP being most prevalent in healthy plaque but significantly less so in abscesses, while FNA shows a higher prevalence in abscesses (13). FNN, once thought to dominate in oral inflammatory sites, has recently been found to be the least prevalent subspecies (13, 17). Notably, FNV often coexists with FNP in healthy mouths and is typically considered benign. Additionally, FNP has been linked to pregnancy complications and oral mucosal diseases (18, 19), while clade 2 of FNA is associated with colon-related disorders (20). In addition to subspecies differences, there are significant variations among strains within the same subspecies. For example, within FNN, the strain ATCC 23726 is a short rod-shaped bacterium, while the clinical strain CTI-2 is filamentous. ATCC 23726 exhibits high transformation capability, but CTI-2’s efficiency is relatively low (21). This strain-specific variability highlights the need for more comprehensive studies beyond the model strain ATCC 23726.

The ability to perform gene deletion in *F. nucleatum* has significantly advanced our understanding of fusobacterial pathogenesis. The current method for gene deletion in *F. nucleatum* relies on a non-replicating “suicide” plasmid that integrates into the chromosome at the target site via homologous recombination (4, 18, 22–25). The success of this method requires the host strain’s high transformation efficiency, which is generally low in most *F. nucleatum* strains. For instance, using 1 µg of shuttle plasmid pCWU6 for transformation, ATCC 23726 can generate more than 10^4 transformants, while most other strains produce fewer than 30 colonies. Not all introduced non-replicating deletion plasmids could successfully integrate into the host chromosome via recombination, with *F. nucleatum* showing a recombination efficiency of around 0.05% (25). Consequently, achieving a high transformation rate is crucial for any meaningful recombination. This has led to ATCC 23726 becoming the model organism for fusobacterial studies, but it does not represent the diversity within the species. The low transformation efficiency in genetically recalcitrant strains is often due to their complex and diverse restriction-modification systems (26). Although methods such as in vitro or in vivo methylation of deletion plasmids have improved transformation efficiency in certain strains like ATCC 25586, these methods are strain-specific, time-consuming, and not broadly applicable (26).

To address these challenges, researchers have introduced temperature-sensitive (Ts) plasmids in other bacteria (27, 28). A replicating plasmid requires a replication protein (RepA) for its maintenance and duplication. These Ts plasmids regulate the activity of the RepA protein at different temperatures, determining whether the plasmid remains independent or integrates into the chromosome. While effective, this approach is impractical for *F. nucleatum*, a strictly anaerobic bacterium, as it would ideally require two anaerobic chambers—one for high temperature and one for low temperature—which is not feasible in our lab.

In seeking alternative methods, we focused on controlling the expression level of RepA rather than its activity. In this study, we developed a conditional plasmid system where an inducible promoter system, consisting of the *fdx* promoter and a theophylline-responsive riboswitch, regulates RepA expression. This system allows the plasmid to replicate within host cells upon induction. Once the inducer is withdrawn, replication ceases, forcing the plasmid to integrate into the chromosome via homologous recombination. We successfully employed this system to create markerless gene deletions in FNN strains (ATCC 23726, CTI-2), an FNP strain (ATCC 10953), an FNA strain (21_1A), and *F. periodonticum* (ATCC 33693).

## RESULTS

### Construction of an inducible system-controlled conditional plasmid for *F. nucleatum*

pCWU6 is an *E. coli/F. nucleatum* shuttle plasmid widely used for expressing both endogenous and exogenous proteins in *F. nucleatum* (29). The replication and maintenance of pCWU6 in the host cells rely on the plasmid replication protein RepA, which is essential for initiating plasmid replication by binding to the origin of replication (*ori_FN_*) located upstream of the *repA* gene (Fig. 1A) (30). Without RepA expression, the plasmid cannot replicate, rendering it non-replicative. This suggests that by controlling *repA* expression, pCWU6 can be theoretically switched from a self-replicating plasmid state to a suicide, non-replicative one within the host.

**Figure 1:**
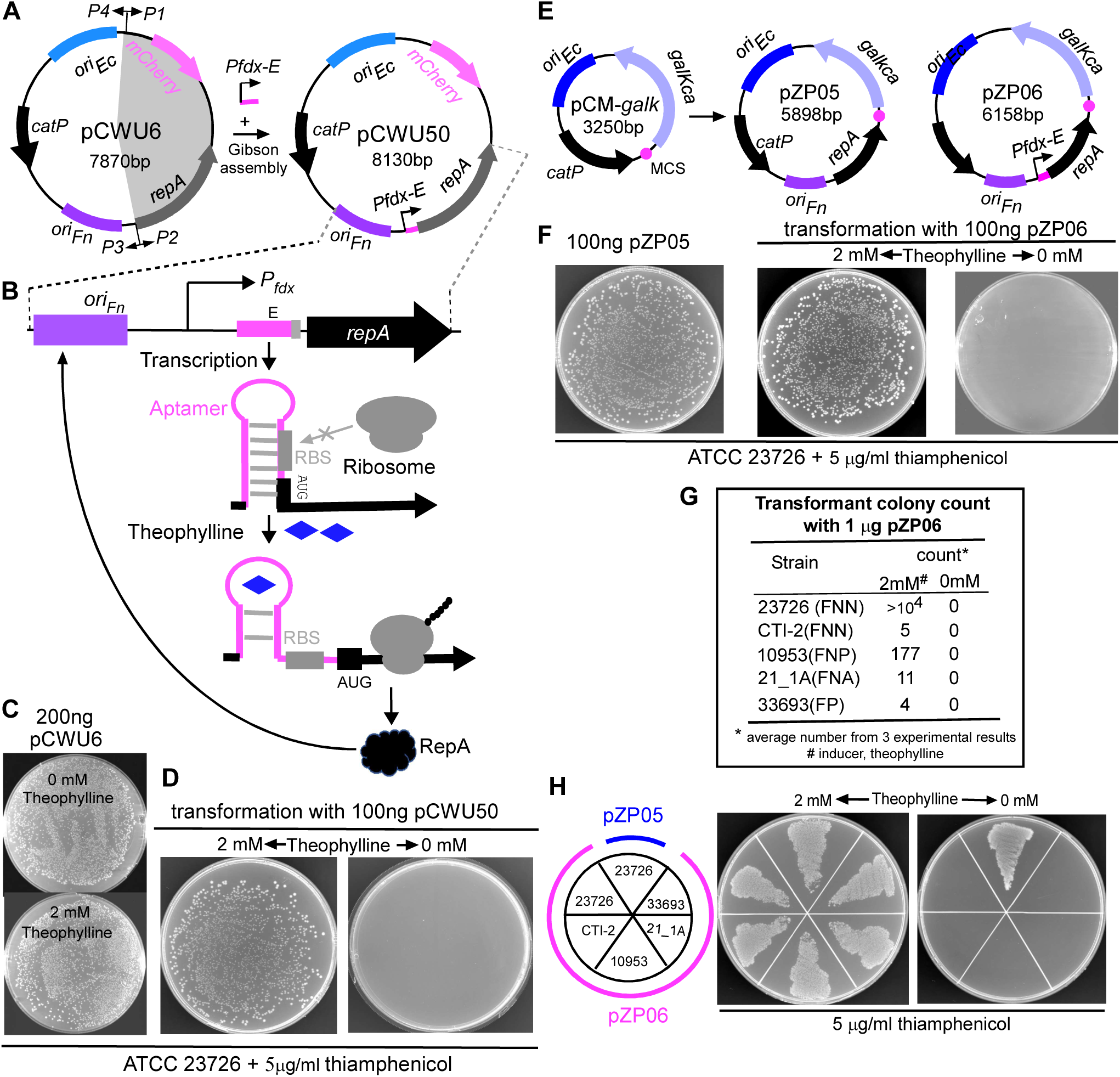
Construction of a riboswitch-controlled conditional plasmid in *Fusobacterium nucleatum*. **(A)** Schematic representation of pCWU50 construction. The P*fdx-E* inducible system was cloned into pCWU6 to drive *repA* expression using Gibson assembly. Primers used to amplify pCWU6 (P1-P4) and key genetic elements such as the origin of replication (*oriFn*), *repA*, and *catP* are indicated. *oriEc* allows plasmid replication in *E. coli*, while *oriFn* and *repA* are required for plasmid replication in *F. nucleatum*. **(B)** Diagram illustrating the theophylline-responsive riboswitch mechanism. Transcription under the P*fdx* promoter is constitutive, but translation is inhibited unless theophylline binds to the riboswitch, causing a conformational change that exposes the ribosome binding site (RBS) and initiates *repA* translation. **(C)** Transformation of pCWU6 into *F. nucleatum* ATCC 23726 is unaffected by the presence or absence of theophylline. 200 ng of pCWU6 was introduced into competent cells, and cells were plated on agar with or without 2 mM theophylline following a 3-hour recovery in an anaerobic chamber. **(D)** The transformation efficiency of pCWU50 in *F. nucleatum* ATCC 23726 is theophylline-dependent. Numerous colonies appeared on thiamphenicol-containing plates with 2 mM theophylline, but none in its absence. Representative results from three independent experiments are shown. **(E)** Construction of pZP05 and pZP06. The *oriFn-repA* fragment from pCWU6 was cloned into pCM-*galK* to create pZP05, and the *oriFn-Pfdx-E-repA* fragment from pCWU50 was cloned into pCM-*galK* to generate pZP06. **(F)** Functional comparison of pZP05 and pZP06. pZP05 behaves similarly to pCWU6, producing numerous colonies when introduced into ATCC 23726. pZP06 behaves like pCWU50, with transformants appearing only in the presence of theophylline. **(G)** Transformation efficiency of pZP06 across various *Fusobacterium* strains. Competent cells from different strains were transformed with pZP06 and plated on thiamphenicol-containing plates with 2 mM theophylline. Colony counts represent the average number from three experimental repeats. **(H)** Growth of strains harboring pZP06 on plates with and without theophylline. Cultures containing pZP06 or pZP05 were grown overnight with 1 mM theophylline, washed, resuspended in fresh TSPC medium, and streaked on plates with or without 2 mM theophylline. Strains with pZP06 died without theophylline but survived with it, while pZP05 showed no dependence on the inducer.

To achieve this control, we constructed the plasmid pCWU50, in which *repA* expression is regulated by the inducible P*fdx-E* system (Fig. 1B). This system consists of the *fdx* promoter, derived from the *Clostridioides sporogenes* ferredoxin gene Clspo_c0087, and a theophylline-responsive riboswitch E unit (31). The riboswitch E unit includes an aptamer and a synthetic ribosome binding site (RBS), which controls gene expression at the translation level (32). We have successfully used this inducible system to regulate gene expression in *F. nucleatum* (33). This system works as follows: transcription under the *fdx* promoter is constitutive during cell growth, but the RBS downstream of the aptamer is sequestered by pairing with the riboswitch stem, preventing translation (Fig. 1B). When theophylline is added, it binds to the riboswitch, causing a conformational change that exposes the RBS to the translation machinery, thereby initiating translation of the *repA* gene from the start codon (AUG) when the ribosome binds to the RBS (Fig. 1B). To validate this system, we used the model organism ATCC 23726 and transformed pCWU50 into its competent cells via electroporation, followed by a 3-hour recovery period in an anaerobic chamber. The recovered cells were then plated onto TSPC agar plates containing thiamphenicol to select for transformants. We expected that if the recovery medium and selection plates lacked theophylline, RepA would not be expressed, leading to a failure in plasmid replication, and no transformants would survive on thiamphenicol-containing agar plates. Conversely, in the presence of theophylline in both the recovery medium and selection plates, RepA would be expressed, enabling plasmid replication and resulting in the appearance of numerous transformants. As expected, innumerable transformants appeared on the plates when pCWU50 was electroporated into ATCC 23726 competent cells with 2 mM theophylline in the recovery medium and selection plates (Fig. 1D, left panel). However, no colonies were observed on plates without the inducer (Fig. 1D, right panel). As a control, the transformation efficiency of pCWU6 in ATCC 23726 was unaffected by the presence or absence of theophylline since *repA* expression in pCWU6 is driven by its native promoter (Fig. 1C).

Due to its large size, pCWU50 posted significant challenges for subcloning, prompting us to develop pZP06, a smaller and more manageable plasmid (Fig. 1E). Before creating pZP06, we first constructed pZP05 by cloning the *oriFN-repA* fragment from pCWU6 into the smaller suicide plasmid pCM-galK, which we had previously used for gene deletion in *F. nucleatum* (29). pZP05 successfully replicated like pCWU6, confirming that *oriFN-repA* was sufficient for replication (Fig. 1F). We then created pZP06 by cloning the *oriFN-Pfdx-E-repA* fragment from pCWU50 into pCM-*galK* (Fig.1E, left panel). As anticipated, pZP06 mimicked pCWU50’s behavior, with transformants appearing only in the presence of theophylline but none in its absence (Fig. 1F, the right two panels). We retained the *galK* gene (*galKca*) from pCM-*galK* in pZP06 to use as a counterselection maker for later markerless gene deletion (Fig. 1E).

The primary goal of creating pZP06 was to facilitate gene deletion in non-model strains with low transformation efficiency. To accomplish this, an essential prerequisite was that this conditional plasmid could be successfully transformed into these genetically recalcitrant strains. To evaluate this, we transformed pZP06 with various fusobacterial strains, including FNN clinical strain CTI-2 (34), FNP strain ATCC 10953, FNA strain 21_1A, and *F. periodonticum* (35) strain ATCC 36693. Despite their lower transformation efficiency, we could always obtain at least one transformant per strain in the presence of theophylline but none without it (Fig. 1G). The transformed cells for each strain grew well with the inducer (Fig. 1H, middle). Still, they died without it (Fig. 1H, right). As a control, the inducer did not affect the growth of the cell with pZP05. These results indicated that P*fdx-E*-controlled *repA* expression effectively switches the plasmid from a replicating to a non-replicating state across various genetic backgrounds.

### Use of the conditional plasmid to delete *galK* genes in non-model fusobacterial strains

To achieve targeted gene deletion, the deletion plasmid must integrate into the chromosome through homologous recombination. Our next step was to determine whether the conditional plasmid could integrate through homologous recombination when the inducer was removed.

To test this, we constructed the plasmid pZP06Δ*galK*, which contains two homologous DNA sequences, each approximately 1 kb in length, derived from the upstream and downstream regions of the *galK* gene (Fig. 2A). The *galK* gene encodes galactokinase, an enzyme critical for galactose metabolism, responsible for phosphorylating D-galactose to galactose-1-phosphate. We chose the *galK* gene for deletion because mutants lacking this gene are easy to screen when D-galactose analog 2-deoxy-D-galactose (2-DG) is included in the selection plates (29). The principle is that 2-DG can be taken up by fusobacterial cells; once inside, it is phosphorylated by galactokinase, producing 2-deoxygalactose-1-phosphate, a toxic compound that inhibits cell growth. As a result, only mutants lacking a functional *galK* gene can survive on plates containing 2-DG, while wild-type strains and those with the integrated *galK* deletion plasmid will be eliminated (Fig. 2G).

**Figure 2:**
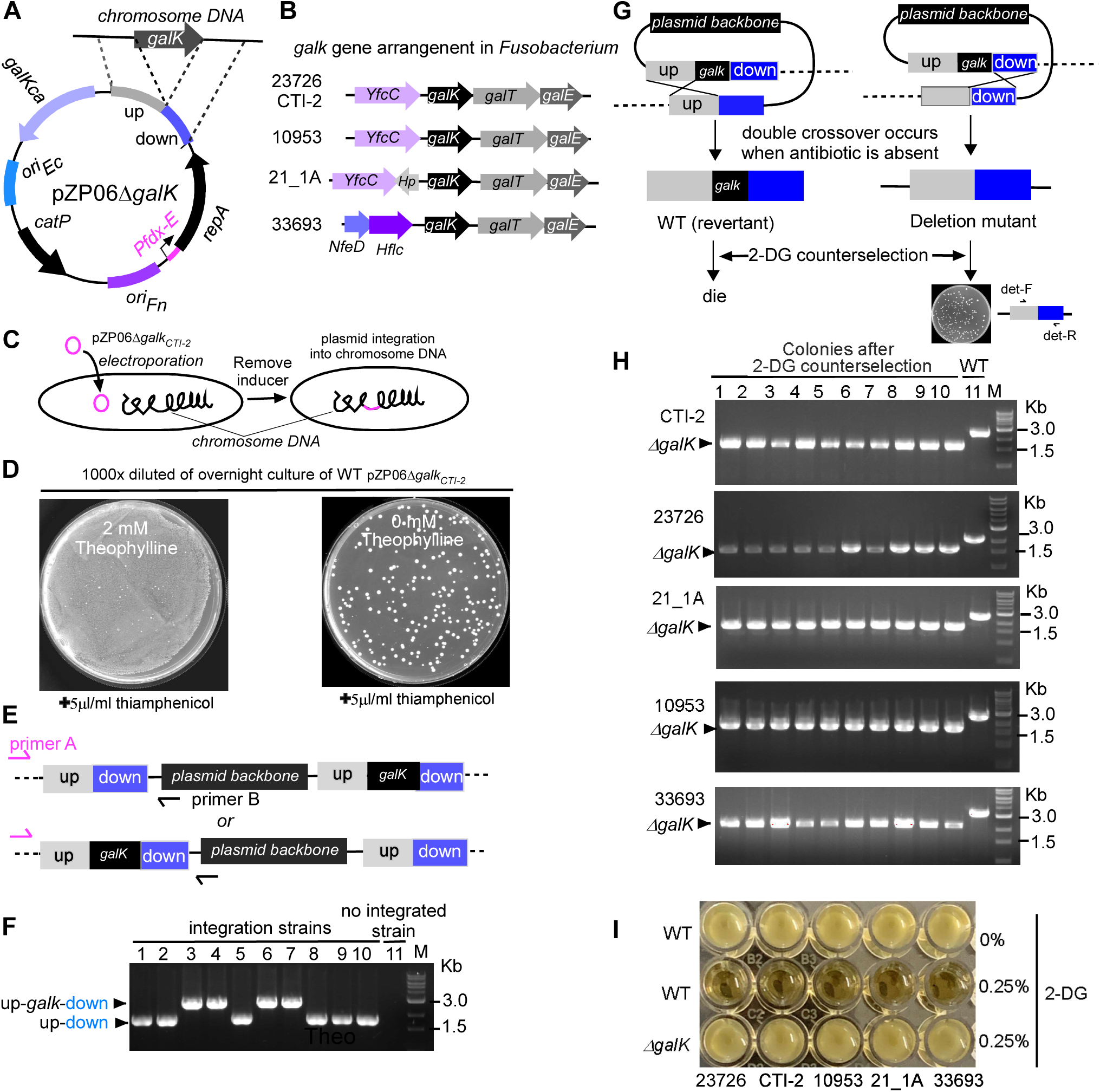
Use of the conditional plasmid pZP06Δ*galK* for targeted deletion of *galK* genes in non-model *Fusobacterium* strains. **(A)** Schematic representation of the pZP06Δ*galK* plasmid, designed to delete the *galK* gene through homologous recombination. The plasmid contains two approximately 1 kb homologous regions derived from the upstream and downstream sequences of the *galK* gene. **(B)** Arrangement of the *galK* gene and its neighboring genes in various *Fusobacterium* strains, illustrating the differences in gene order and orientation. **(C)** Transformation and integration strategy for pZP06Δ*galK* in strain CTI-2. Competent CTI-2 cells were transformed with pZP06Δ*galK* in the presence of theophylline, and transformants were selected on thiamphenicol-containing plates. Cells were subsequently incubated without the inducer to facilitate plasmid integration into the chromosome through homologous recombination. **(D)** Growth of CTI-2 cells transformed with pZP06Δ*galK* on TSPC agar plates with or without theophylline in the presence of 5 µg/ml thiamphenicol. Approximately 100 colonies appeared on plates without the inducer, while a bacterial lawn was observed on plates with theophylline, indicating successful plasmid integration. **(E)** PCR analysis to confirm plasmid integration. Primer A, located upstream of the *galK* homologous region, and Primer B, annealing to the plasmid backbone, were used to amplify products from the transformed cells. Distinct PCR products indicated successful integration at either the upstream or downstream homologous region. **(F)** PCR results show upstream integration (1.8 kb product) in six colonies and downstream integration (3.0 kb product) in four colonies. Non-integrated strains yielded no PCR products. **(G)** Strategy for isolating *galK* mutants in strain CTI-2. Following the first crossover event, a second crossover event is induced in the absence of antibiotics and theophylline, leading to the excision of the plasmid and resulting in either an in-frame deletion mutation or reversion to the wild-type genotype. Screening for *galK* mutants using 2-DG counterselection. Cells with a functional *galK* gene (wild-type or plasmid-integrated) are eliminated, while *galK* mutants survive. **(H)** PCR analysis of *galK* mutants using primers det-F/R (see Fig. 2G) in five *Fusobacterium* strains (CTI-2, ATCC 23726, 21_1A, ATCC 10953, and *F. periodonticum* ATCC 333693) selected on 2-DG counterselection plates. PCR confirmed all 10 randomly picked colonies from each strain as *galK* mutants. **(I)** Successful generation of *galK* mutants in multiple *Fusobacterium* strains (ATCC 10953, ATCC 23726, 21_1A, and *F. periodonticum* ATCC 333693) using the pZP06ΔgalK strategy. All mutants demonstrated sensitivity to 2-DG, validating the effectiveness of the conditional plasmid system.

Each fusobacterial strain tested harbors a single copy of the *galK* gene, but the gene’s arrangement varies across strains, particularly in its position relative to the downstream *galT* gene and the differences in adjacent upstream genes (Fig. 2B). To accommodate these variations, we designed strain-specific *galK* deletion plasmids, such as pZP06-Δ*galK*_CTI-2._ To demonstrate the procedure for gene deletion, we used the plasmid tailored for strain CTI-2 as an example. This plasmid was introduced into competent CTI-2 cells via electroporation in the presence of theophylline (Fig. 2C). Twelve transformants were obtained, and one was selected for further analysis. This transformant (CTI-2 pZP06-Δ*galK_CTI-2_*) was grown overnight in TSPC broth with 2 mM theophylline. After harvesting, the cells were washed and resuspended in inducer-free TSPC broth for a 7-hour incubation to deplete residual inducer (Fig. 2C). The culture was then serially diluted and plated on TSPC agar containing 5 µg/ml thiamphenicol. We hypothesized that in the absence of the inducer, pZP06-Δ *galK_CTI-2_* would shift from a replicative to a non-replicative suicide plasmid, and under antibiotic selection, a small portion of the plasmids would integrate into the chromosome through a low frequency of homologous recombination.

As expected, approximately 100 colonies appeared on the plates without the inducer, while a bacterial lawn was observed on plates with the inducer (Fig. 2D). To confirm plasmid integration, we designed two primers: Primer A, which anneals a region just before the upstream homologous arm of the *galK* gene in the chromosome, and Primer B, which anneals to the plasmid backbone (Fig. 2E). Without plasmid integration, no PCR product was expected. However, if integration occurred, distinct PCR products of different sizes would be observed depending on the recombination site: a smaller product for upstream integration and a larger product for downstream integration encompassing the *galK* deletion region (Fig. 2E). PCR analysis of ten randomly selected colonies from the no-inducer plates confirmed our predictions. Six colonies produced a 1.8 kb PCR product, indicative of upstream integration. In contrast, four produced a 3.0 kb PCR product, indicative of downstream integration (Fig.2F). No PCR products were obtained from non-integrated strains (Fig. 2F, the last sample lane), validating that the conditional plasmid integrates into the chromosome through a single crossover event at one of the homologous regions in the absence of the inducer.

To isolate the *galK* mutant in strain CTI-2, we propagated one of the integrated clones without antibiotics and inducer, allowing a second crossover event between the duplicated homologous regions. This event leads to plasmid excision from the chromosome (Fig. 2G). Given that the plasmid contains two homologous fragments of similar size, the secondary recombination event has an equal probability of occurring at either site, potentially resulting in an in-frame deletion mutation or reversion to the wild-type genotype (Fig. 2G). However, due to the infrequency of double crossover events, most cells in the culture remained plasmid-integrated even without antibiotic selection. At this stage, the advantage of choosing *galK* for deletion became evident. By plating the culture on TSPC plates containing 2-DG, we could quickly screen for mutants, as cells with a functional *galK* gene, including plasmid-integrated strains or revertant wild types, would die, while only *galK* mutants would survive. As anticipated, colony PCR confirmed all 10 colonies randomly picked from the 2-DG plates as *galK* mutants (Fig. 2H).

Using this strategy, we also successfully generated *galK* mutants in strains ATCC 10953, ATCC 23726, 21_1A, and *F. periodonticum* ATCC 333693 (Fig. 2H). All these *galK* mutants were sensitive to 2-DG (Fig. 2I), demonstrating the effectiveness of this conditional plasmid system in generating gene deletions in genetically recalcitrant strains.

### Use of the conditional plasmid with *galK* as a counterselection marker to make gene deletion in *F. periodontium*

In constructing pZP06-Δ*galK*, we retained the *galKca* gene from pCM-*galK*, even though it was not needed for the plasmid’s function. This *galK* gene, derived from *Clostridium acetobutylicum* ATCC 824, has been successfully used as a counterselection marker in *Clostridium perfringens* and *F. nucleatum* ATCC 23726 (29, 36), but its efficacy in other non-model fusobacterial strains had not been tested. To use *galK* for counterselection, a *galK*-minus recipient strain was necessary(29). Having generated *galK* mutants for various non-model strains, we tested whether this conditional plasmid, combined with the *galK* counterselection, could create markerless in-frame deletions.

We selected *F. periodontium* Δ*galK* as the recipient strain and constructed a pZP06-based allelic exchange vector, pZP06Δ*luxS*, to delete the *luxS* gene (FperA3_010100009561, https://img.jgi.doe.gov/) (Fig. 3A). This gene encodes S-ribosyl homocysteine lyase, an enzyme that produces the quorum-sensing molecule autoinducer-2 (AI-2), which bacteria use to coordinate group behaviors based on population density (37). Using the *Vibrio harveyi* BB170 reporter strain, which induces luminescence in response to AI-2 (38), we confirmed that *F. periodontium* ATCC 33693 produces AI-2 at levels comparable to those made by AI-2-producing *E. coli* (Fig. 3E). In contrast, only background signals were detected in FNN strains ATCC 23726 and CTI-2, which lack the *luxS* gene (Fig. 3E, last two columns).

**Figure 3:**
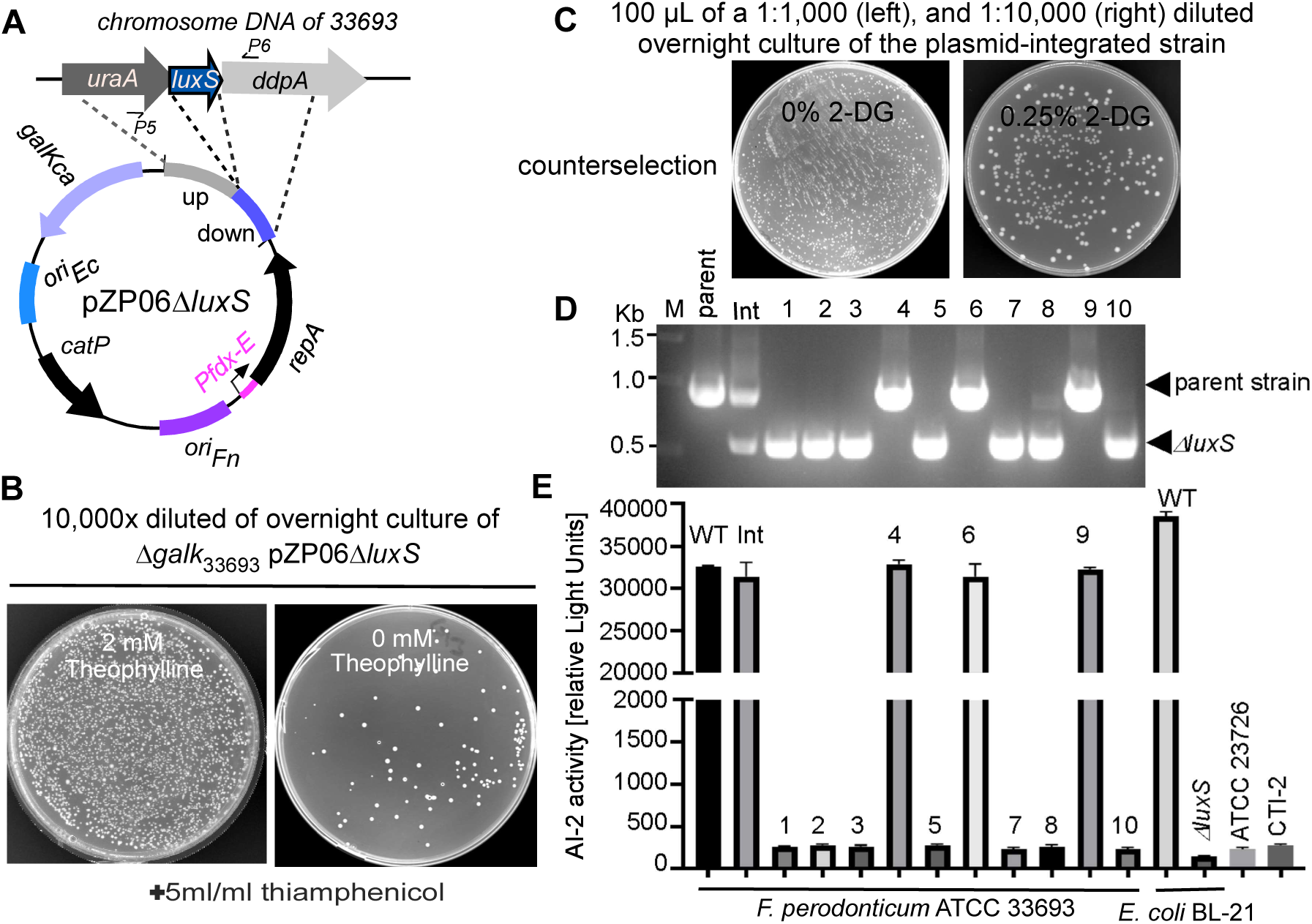
Application of pZP06Δ*luxS* with *galK* as a counterselection marker for gene deletion in *Fusobacterium periodonticum*. **(A)** Schematic of the pZP06Δ*luxS* allelic exchange vector designed to delete the *luxS* gene in *F. periodonticum* Δ*galK*. The *luxS* gene encodes S-ribosyl homocysteine lyase, essential for producing the quorum-sensing molecule autoinducer-2 (AI-2). **(B)** Selection of plasmid-integrated *F. periodonticum* Δ*galK* strains after transformation with pZP06Δ*luxS*. Transformed cultures were plated on thiamphenicol-containing plates with or without 2 mM theophylline. The left panel shows a bacterial lawn in the presence of theophylline, indicating no plasmid integration. The right panel shows isolated colonies without theophylline, where plasmid integration into the chromosome occurred via homologous recombination. **(C)** Screening for plasmid excision using 2-DG counterselection. The second crossover event, induced by growing the culture in an antibiotic-free medium, led to a few 2-DG-resistant colonies, indicating successful plasmid loss and potential in-frame deletion of *luxS*. **(D)** PCR analysis of ten 2-DG-resistant colonies with primers P5/P6 (indicated in Fig. 3A) revealed seven were in-frame *luxS* deletion mutants, while the remaining three retained the parent strain’s genotype. **(E)** AI-2 bioassay results. *F. periodonticum* ATCC 33693 produced AI-2 at levels similar to AI-2-producing *E. coli*, as measured by luminescence in the *Vibrio harveyi* BB170 reporter strain (see Methods for details). In contrast, *luxS* deletion mutants showed significantly reduced luminescence, confirming the loss of AI-2 production.

We introduced pZP06Δ*luxS* into *F. periodontium* Δ*galK* via electroporation and selected transformants on plates containing 2 mM inducer. Of the five transformants obtained, one was chosen for further study. This transformant was grown overnight in TSPC broth with inducers, followed by washing and resuspension in inducer-free TSPC broth before plating on TSPC agar with 5 µg/ml thiamphenicol to screen for colonies where the plasmid had integrated into the chromosome. Any colonies that appeared were considered pZP06Δ*luxS*-integrated strains (Fig. 3B).

One plasmid-integrated colony was randomly chosen and grown overnight in a TSPC medium with antibiotics. The following day, the culture was diluted 1:1,000 in a fresh TSPC medium without antibiotics to allow the second crossover event, which would result in an in-frame deletion of *luxS*. The culture was grown to an OD_600_ of about 1.0, then diluted 1:10,000 and plated on TSPC agar containing 2-DG. A control plate without 2-DG was used to estimate the frequency of plasmid excision. As shown in Figure 3C, 2-DG counterselection allowed only a tiny fraction of cells to survive. Based on the ratio of 2-DG-resistant cells to total cells, we estimated that approximately 1 in 2000 cells underwent the second recombination event, resulting in plasmid loss.

We randomly selected ten 2-DG-resistant colonies and tested them for antibiotic sensitivity to confirm plasmid loss. Colony PCR analysis showed seven colonies were in-frame deletion mutants, while the remaining three retained the parent strain’s genotype (Fig. 3D). Deletion of the *luxS* gene abolishes AI-2 production. As expected, the AI-2 assay showed that all *luxS* mutant colonies exhibited the lowest luminescence (Fig. 3E). In contrast, colonies with the same genotype as the parent strain displayed high luminescence levels similar to those of the parent and plasmid-integrated strains, confirming the functionality of the AI-2/LuxS quorum sensing system in *F. periodontium* (Fig. 3E). The successful deletion of the *luxS* gene in *F. periodontium* demonstrates that this conditional plasmid system can effectively create markerless gene deletions in non-model organisms using *galK* as a counterselection marker.

### Use of the conditional plasmid with *sacB* as a counterselection marker to make gene deletion in non-model strains

While the *galK*-based counterselection method has proven highly effective for unmarked deletion mutagenesis, it is limited by the requirement for a host strain lacking the *galK* gene. To overcome this limitation and extend the applicability of gene deletion strategies to wild-type backgrounds, we recently developed a *sacB*-based counterselection system for *F. nucleatum* ATCC 23726 (39). Our last goal was to determine whether this conditional plasmid system, combined with *sacB* as a counterselection marker, could facilitate gene deletions in non-model strains.

To test this, we constructed the plasmid pZP06B, where the *galK* gene and its promoter from pZP06 were replaced with *sacB* and the *rpsJ* promoter from pZP02 (39) (Fig. 4A). To streamline the cloning process, we introduced pZP06C, which includes a mCherry reporter cassette (P*rpsJCd-mCherry*) in the multiple cloning site (MCS) regions of pZP06B (Fig. 4B). The mCherry cassette provided a visual marker, enabling easy identification of positive clones— colonies containing pZP06C glowed red, simplifying the selection process (21).

**Figure 4:**
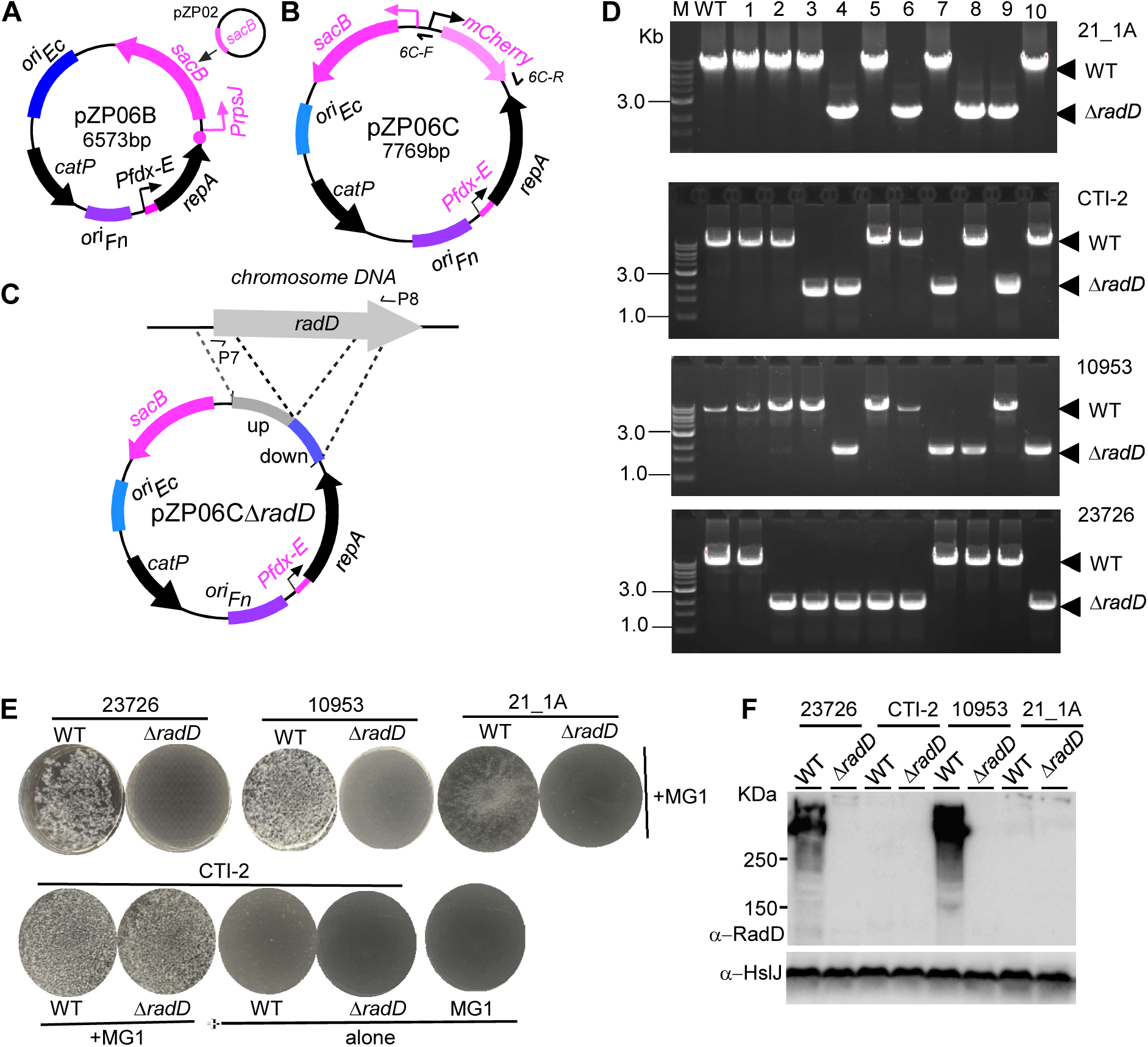
Application of the conditional plasmid with *sacB* as a counterselection marker for gene deletion in non-model *Fusobacterium* strains. **(A)** Schematic of the pZP06B plasmid, in which the *galK* gene was replaced with *sacB* under the control of the *rpsJ* promoter, enabling *sacB*-based counterselection. **(B)** Schematic of pZP06C, an enhanced version of pZP06B, containing a *mCherry* reporter cassette in the multiple cloning site (MCS) to provide a visual marker for identifying positive clones. **(C)** Construction of pZP06CΔradD for deleting the *radD* gene in *Fusobacterium nucleatum* strain 21_1A. The *radD* gene encodes a large surface adhesion protein that facilitates coaggregation with oral bacteria. **(D)** PCR analysis using primer pair P7/P8 (indicated in panel C) of ten sucrose-resistant colonies after *sacB* counterselection in four *Fusobacterium* strains (21_1A, CTI-2, ATCC 10953, and ATCC 23726). In strain 21_1A, four colonies produced a 2.0 kb amplicon, indicating a *radD* deletion, while the remaining six colonies produced the 8.3 kb wild-type amplicon. In the other strains, the PCR results showed varying numbers of mutants and wild-type colonies, with smaller amplicons corresponding to the *radD* deletions and larger amplicons corresponding to the wild-type genotype. **(E)** Coaggregation assays with *A. oris* MG-1. Deletion of the *radD* gene abolished coaggregation in strains 21_1A, ATCC 10953, and ATCC 23726 but not in strain CTI-2. The wild-type and ΔradD mutant cells for each strain were cultured in TSPC medium to the stationary phase, then collected, washed, resuspended in coaggregation buffer, and assessed for coaggregation with an equal amount of *A. oris* cells. Representative results are shown from repeated experiments with all screened mutant colonies. **(F)** Western blot analysis of RadD expression in wild-type and ΔradD mutant strains. Cells used in the coaggregation assays in panel E underwent SDS-PAGE, followed by immunoblotting. Antibodies specific to RadD (α-RadD) and HsIJ (α-HsIJ) were used for detection, with HsIJ serving as a loading control (29). Molecular weight markers (in kilodaltons) are indicated on the left side of the blot.

We then applied this system to create pZP06CΔ*radD*, a plasmid to delete the *radD* gene (Fig. 4C). The *radD* gene encodes a large surface adhesion protein RadD that facilitates *F. nucleatum*’s coaggregation with certain oral bacteria, particularly Gram-positive early colonizers, critical for dental plaque formation (4). While *F. periodontium* lacks a *radD* homolog, most *F. nucleatum* strains, including ATCC 23726, CTI-2, ATCC 10953, and 21_1A, possess a copy of the *radD* gene, with sequence identities ranging from 66.7% to 82.7% (Fig. S1). Due to this variability, we constructed four versions of pZP06CΔ*radD* tailored to each strain.

To illustrate this process, we describe the deletion of the *radD* gene in FNA strain 21_1A using this new gene deletion system. The plasmid pZP06CΔ*radD*_21_1A_ was introduced into competent cells of strain 21_1A. In the presence of the inducer, nine transformants were obtained in one electroporation reaction, one of which was selected for further analysis. This transformant was grown in a medium with antibiotics but without the inducer, causing the plasmid pZP06CΔ*radD*_21_1A_ to integrate into the chromosome, resulting in a plasmid-integrated strain. One plasmid-integrated strain was selected for *sacB* counterselection by growing it on a sucrose-containing TSPC medium. After 4.5 days, ten randomly selected sucrose-resistant colonies were tested for antibiotic sensitivity to screen for potential *radD* deletion mutants. Colony PCR using the primer pair Δ*radD*-F/R was then performed to analyze their genotypes. Figure 4D shows four of the ten colonies produced amplicons of approximately 2.0 kb, indicating a gene deletion. The remaining six colonies displayed the expected wild-type amplicons of approximately 8.3 kb (Fig. 4D).

Using this approach, we successfully generated *radD* mutants in the non-model strains CTI-2 and ATCC 10953 and the model strain ATCC 23726 (Fig. 4D). We further validated the *radD* mutants by conducting a coaggregation assay for each strain. For strains 21_1A, ATCC 10953, and ATCC 23726, deletion of the *radD* gene abolished fusobacterial coaggregation with *Actinomyces oris* MG-1 (Fig. 4E). Interestingly, the CTI-2 Δ*radD* mutant retained its coaggregation ability, which was unexpected, as we believed that only the RadD protein in *F. nucleatum* was responsible for coaggregation with MG-1 (Fig. 4E). To investigate this further, we performed Western blot analysis to detect RadD expression in both wild-type and Δ*radD* mutant strains. Consistent with the coaggregation results, ATCC 23726 and 10953 exhibited high levels of RadD expression in the wild-type background. Still, none in the mutant cells (Fig. 4F). The antibody against RadD was generated using a recombinant RadD protein with a highly conserved region between the 44th and 200th amino acids in ATCC 23276’s RadD (Fig. S1). This should theoretically allow the antibody to recognize any RadD protein. Surprisingly, however, no RadD signal was detected in either the wild-type or Δ*radD* mutant strains of CTI-2 and 21_1A (Fig. 4F). Despite these unexpected findings, our results demonstrate that using a conditional plasmid with *sacB* as a counterselection marker is feasible and effective for achieving gene deletions in the wild-type backgrounds of non-model strains. This approach expands the toolkit for genetic manipulation in *F. nucleatum*, enabling further exploration of this pathogen’s diverse behaviors and virulence factors.

## DISCUSSION

*F. nucleatum* is characterized by considerable genetic and phenotypic diversity across its various strains, which has historically posed challenges for genetic manipulation, particularly in strains with low transformation efficiency (26, 33). This limitation has significantly restricted our understanding of the full spectrum of *F. nucleatum* biology and its pathogenic potential. To address this, we developed a conditional plasmid system that leverages a riboswitch-based inducible system to control the expression of the plasmid’s replication protein, RepA. This innovation allows the plasmid to replicate in the presence of an inducer and integrate into the host chromosome via homologous recombination when the inducer is removed. Combining this system with *galK* or *sacB* as counter-selection markers enabled unmarked gene deletions in genetically recalcitrant strains of *F. nucleatum*.

Our approach to constructing this conditional plasmid differs fundamentally from traditional methods that utilize temperature-sensitive (Ts) plasmids. The latter typically relies on error-prone PCR to screen for a mutated RepA protein active only at lower temperatures but nonfunctional at higher temperatures, thereby controlling the plasmid’s replication and integration into the chromosome by altering the growth temperature (28, 40). While effective in many Gram-positive and Gram-negative bacteria, this method is impractical for strict anaerobes like *F. nucleatum*, as it would ideally necessitate the use of two anaerobic chambers set to different temperatures—an infeasible solution for most laboratories. Additionally, Ts plasmid-based methods often make the gene deletion process time-consuming due to the slow growth of bacteria at lower temperatures. We found some strains of *F. nucleatum* cannot form colonies in temperatures below 30°C for two weeks. In contrast, our conditional plasmid system uses theophylline as an inducer, which does not affect bacterial growth (32). Although we tested a limited number of fusobacterial strains, we successfully achieved gene deletions in each, demonstrating the broad applicability of our system. This method can be extended to other *F. nucleatum* strains and related bacteria. Interestingly, as we were finalizing this manuscript, a similar system using a theophylline-responsive riboswitch to control RepA expression was successfully employed in *Clostridium difficile* (41), further validating our approach.

The critical factor in using this conditional plasmid for gene deletion in hard-to-transform fusobacterial strains is the ability to transform the deletion plasmid into the cells. Our standard procedure involves extracting the deletion plasmid and introducing it into competent cells of the host strain via electroporation (42). The quality of the competent cells is crucial; cells from the stationary phase often have a higher transformation efficiency than those in the logarithmic phase. Using this method, we could consistently obtain transformants if we used a high concentration of competent cells and a sufficient amount of the pZP06C-based deletion plasmid, ideally around 1 µg per electroporation reaction. Occasionally, a single electroporation may not yield a transformant, necessitating multiple attempts. With this approach, we successfully transformed the pZP06C into strains historically challenging to transform, such as ATCC 25586 and ATCC 51191, belonging to the FNN and FNA subspecies, respectively.

Among the three non-model *Fusobacterium nucleatum* strains and one *F. periodontium* strain we tested, FNP ATCC 10953 exhibited relatively higher transformation efficiency (Fig. 1G). This efficiency helps explain why gene deletions have been achieved in this strain. However, it’s important to note that, despite this transformation capability, only two marked gene deletions— *radD* and *fad-I* (43–45)—have been successfully performed so far. While transformation is feasible, extensive gene deletion work in this strain remains challenging and somewhat limited. To our knowledge, no gene deletion has been reported in any FNA strain. We are the first to create a gene deletion in the FNA strain 21_1A, a clinical isolate recovered from human colon cancer samples (46). Recent years have seen accumulating evidence that FNA strains are enriched in human colorectal cancer (CRC) tumors and are associated with cancer development (47, 48). Interestingly, studies have suggested that within the FNA subspecies, there are two distinct clades, C1 and C2, with only FNA C2 dominating the CRC tumor niche (20). FNA differs significantly from other subspecies, with unique gene loci and phenotypes (11, 13, 20). It is better to use strains from FNA, particularly C2 strains, in animal models to establish the relationship between FNA and CRC development. However, these FNA strains are genetically recalcitrant, so most experiments currently use the FNN strain ATCC 23726 as a substitute (45, 49, 50). Given the many differences between these subspecies, conclusions drawn from such experiments may be problematic. To truly understand *F. nucleatum*’s complex nature and disease-causing capabilities, expanding research to include a broader range of strains is essential, laying the groundwork for better animal study and clinical practices.

RadD is a surface protein that facilitates *F. nucleatum*’s physical interactions with many Gram-positive oral bacteria(4). However, RadD’s function varies across the four subspecies. For example, only FNP’s RadD can bind to *Streptococcus mutans* SpaP, while RadD from other subspecies cannot (43). Recent reports also indicated that RadD from FNN and FNA can bind to Siglec-7, a member of the sialic acid-binding immunoglobulin-like lectins predominantly expressed on NK cells, but not RadD from FNP (51). Differences in the amino acid sequence of RadD among subspecies and even between strains within the same subspecies may explain these functional variations. Deleting the *radD* gene across multiple strains revealed strain-specific differences in coaggregation with *A. oris*, particularly in the CTI-2 strain (Fig. 4E), suggesting the involvement of other adhesins and highlighting the species’ complexity and diversity. Whether RadD is expressed in CTI-2 or simply not detectable by the antibody developed against RadD from ATCC 23726 requires further investigation. In the 21_1A strain, it is clear that the RadD protein was expressed, as deletion of the *radD* gene abolished its coaggregation with *A. oris*. Our inability to detect RadD in the 21_1A strain using a RadD-specific antibody raises intriguing questions about potential variations in the RadD protein across different strains. The antibody against RadD was generated using a recombinant RadD protein with a highly conserved region between the 44th and 200th amino acids in ATCC 23726’s RadD. Compared to RadD in 23726, RadD in CTI-2 has only three amino acid variations, and RadD in 21_1A has two amino acid differences specifically within this alignment region (44th to 200th amino acids) (Fig. S1). Minor amino acid substitutions in some proteins like p53 and the influenza hemagglutinin protein could significantly impact antibody recognition, rendering the protein undetectable via standard Western blot analysis (52, 53). This may be the case here, but further confirmation is required. This observation underscores the importance of exploring molecular differences across *F. nucleatum* strains, as they may contribute to the strain-specific behaviors observed.

Overall, this study introduces a powerful new tool for genetic studies in *F. nucleatum* and related species. The conditional plasmid system we developed represents a significant technical advancement and opens new avenues for exploring the genetic basis of phenotypic diversity and pathogenicity in these organisms. Future research using this system will likely provide deeper insights into the genetic factors contributing to the varied disease potentials among different *F.* nucleatum strains and subspecies.

## MATERIALS AND METHODS

### Bacterial strains and growth conditions

Bacterial strains and plasmids used in this study are listed in Table S1. Fusobacterial species/strains were grown in Tryptic Soy Broth (TSB) supplemented with 1% Bacto Peptone and 0.05% freshly made cysteine (TSPC) or on TSPC agar plates anaerobically (85% N_2_, 10% CO_2_, 5% H_2_) at 37°C, for the selection and counter-selection of transformants, cultures and plates were supplemented with thiamphenicol at 5 µg/ml with and without 2 mM filter-sterilized theophylline. *Escherichia coli* strains were grown in Luria–Bertani (LB) broth (Difco) with aeration at 37°C. *E. coli* strains carrying plasmids were grown in LB media containing 20 µg/ml chloramphenicol (Sigma).

### Plasmids construction

All plasmids used in this study are listed in Table S1, and they were constructed via Gibson Assembly Master Mix kit (New England Biolabs) according to manufacturers’ instructions. In brief, A 2x repliQa HiFi ToughMix from Quantabio (Catalog #: 95200) was used for all PCR reactions. A linearized vector backbone obtained from inverted PCR and inserted PCR fragments were mixed with 2x Gibson Assembly Master Mix in a total volume of 20 μL at 50°C for 20 minutes. Two μL mixture was transformed into *E. coli* competent cells according to transformation protocol. PCR and sequencing confirmed the plasmids. The PCR primers used in this study are listed in Table S2. All plasmids were propagated using *E. coli* DH5α as the cloning strain and electroporated into competent fusobacterial species/strains.

#### (i) pCWU50

To evaluate the ability of the P*fdx-E* inducible system to regulate the expression of *repA* in pCWU6 (29), we cloned the P*fdx-E* promoter region from plasmid pZP4C (33) and inserted it directly upstream of the *repA* start codon. The P*fdx-E* promoter was amplified using the primer pair ribo-E-F and pfdx-E-R. Initially, we attempted to obtain the pCWU6 backbone using an inverse PCR approach with primers P2 and P3 (Fig. 1A); however, this method was unsuccessful. Consequently, we decided to amplify pCWU6 in two separate fragments. The first fragment was amplified with primers P1 and P2, while the second was amplified using primers P3 and P4. The two amplified fragments and the PCR product containing P*fdx-E* were then combined and subjected to a Gibson assembly reaction, resulting in the construction of pCWU50.

#### (ii) pZP05

The purpose of constructing pZP05 was to determine whether the *oriFn-repA* elements are the only genetic components required to replicate pCWU6. To achieve this, the *ori_Fn_-repA* region was amplified from pCWU6 using the primer pair ori-repA-F and ori-repA-R. Simultaneously, the backbone of pCM-*galK* (36) was generated through inverse PCR using the primer pair 2pCm-galK-F and 2pCm-galK-R, which were designed to anneal to a region between the *catP* gene and the multiple cloning site (MCS) in pCM-galK (Fig. 1E). The amplified *ori_Fn_-repA* region was then ligated to the linearized pCM-galK backbone using a Gibson assembly reaction, resulting in the construction of pZP05. The sequence of pZP05 was subsequently confirmed by Sanger sequencing.

#### (iii) pZP06

To construct the smaller conditional plasmid pZP06, the *ori_Fn_-Pfdx-E-repA* region was PCR-amplified from pCWU50 using the primer pair ori-repA-F and ori-repA-R. The amplified DNA fragment was cloned into the linearized pCM-galK backbone via a Gibson assembly reaction, creating pZP06. The integrity and accuracy of pZP06 were validated through whole-plasmid sequencing.

#### (IV) pZP06Δ*galK*

To target the *galK* gene in each of the five tested strains (ATCC 23726, CTI-2, 10953, 21_1A, and ATCC 33693), we constructed five distinct pZP06Δ*galK* deletion plasmids. This was necessary because the upstream and downstream regions of the *galK* gene vary among these strains, requiring the design of a separate plasmid for each one. The construction process began with an inverse PCR using the primer pair pZP06-F/R to linearize the pZP06 plasmid, the cloning backbone. Note that pZP06-F/R are annealed at the region in the MCS of pZP06 (Fig. 1B). The upstream and downstream regions of the *galK* gene were then amplified from the genomic DNA of each strain using strain-specific primer pairs (listed in Table S2). The two PCR amplicons for each strain were fused by overlapping PCR, and the fused fragments were then ligated into the linearized pZP06 backbone via a Gibson assembly reaction. The resulting plasmids, one for each strain listed in Table S1, were confirmed by PCR and sequencing.

#### (V) pZP06Δ*luxS*

To create an in-frame deletion of the *luxS* gene (from amino acid 7 to amino acid 134; FperA3_010100009561, accessible at https://img.jgi.doe.gov/), we amplified 1.0-kb fragments upstream and downstream of *luxS* using PCR with the primer pairs 33693luxSupF/R and 33693luxSdnF/R, respectively. These two PCR amplicons were then fused via overlapping PCR. To clone the fused fragment into the pZP06 plasmid, pZP06 was linearized by inverse PCR using the primer pair pZP06-F/R (Figure 2A). The fused fragment and the linearized pZP06 backbone were subsequently joined using a Gibson assembly reaction. The resulting plasmid, pZP06ΔluxS, was confirmed and validated by PCR and sequencing.

#### (VI) pZP06B and pZP06C

To substitute the *galKcm* gene with *sacB* as a new counterselection marker in pZP06, we used an inverse PCR with the primer pair galKrem-F2/R2 to concurrently remove the *fdx* promoter and *galK* from pZP06, leaving a linearized pZP06 backbone. The *sacB* gene and the rpsJ promoter were amplified from pZP02 using the sacB-F/R primers. The resulting PCR product (P*rpsJ-sacB)* was then ligated with the linearized plasmid backbone through a Gibson assembly reaction, forming pZP06B (Fig. 4A). To streamline the screening of positive clones and enhance the efficiency of Gibson assembly, we developed the pZP06C plasmid. This plasmid includes the *mCherry* gene, which encodes a red fluorescent protein positioned within the multiple cloning site (MCS) regions (Fig. 4A). The *mCherry* gene is flanked by the annealing sites of primers 6C-F and 6C-R, allowing for easy cloning of allelic exchange cassettes. To create pZP06C, we began by linearizing the pZP06B plasmid using an inverse PCR with the primer pair 6C-F/R. Simultaneously, the *mCherry* gene, driven by an *rpsJCd* promoter, was amplified from the pCWU6 template using the mCherry-F/R primer pair. The *mCherry* amplicon was combined with the linearized pZP06B in a Gibson assembly reaction. Bacteria transformed with the resulting pZP06C plasmid produce red colonies, a phenotype resulting from *mCherry* expression.

#### (VII) pZP06CΔ*radD*

Following a procedure similar to the *galK* deletion, the upstream and downstream fragments of the *radD* gene were amplified for each strain using genomic DNA from ATCC 23726, strain CTI-2, ATCC 10953, and 21-1A as templates, along with the corresponding primer pairs (Table S2). The two PCR amplicons were then fused by overlapping PCR. To prepare the backbone for cloning, the *mCherry* gene was removed from pZP06C by PCR using the primer pair 6C-F/R, generating the linearized pZP06C backbone. The fused *radD* fragments and the pZP06C backbone were joined via a Gibson assembly reaction. The resulting series of pZP06CΔ*radD* plasmids, specific to each strain, were confirmed by PCR and sequencing.

### Deletion of *galK* in fusobacterial species/strains

The plasmids pZP06Δ*galK* were introduced into the corresponding fusobacterial competent cells via electroporation (42). The antibiotic-resistant colonies were selected on TSPC agar plates supplemented with 5 µg/mL thiamphenicol and 2 mM theophylline. One to two thiamphenicol-resistant colonies were selected and cultured overnight in TSPC broth containing thiamphenicol and 1 mM theophylline. Following overnight incubation, 1 mL of the culture was pelleted, washed twice with TSPC, and resuspended in 8 mL of inducer-free TSPC broth for continued growth over an additional 7-hour period to eliminate residual inducer. The culture was then diluted 1,000-fold and 10,000-fold and spread onto TSPC agar plates containing 5 µg/mL thiamphenicol to select for plasmid-integrated strains. Theophylline removal silenced the *repA* gene, facilitating plasmid integration into the chromosomal DNA through homologous recombination under antibiotic selection. A thiamphenicol-resistant colony was then inoculated in TSPC broth without antibiotics and cultured overnight. This culture was again diluted 1,000-fold and 10,000-fold into fresh TSPC medium and spread onto TSPC agar plates containing 0.25% 2-DG to screen for cells that had lost the plasmid through a loop-out event. Approximately 10 colonies were re-streaked onto TSPC plates and TSPC plates containing 5 µg/mL thiamphenicol to test for thiamphenicol sensitivity. Thiamphenicol-sensitive strains, indicating plasmid loss, were selected and further analyzed by PCR using the primers det-F/R (Fig. 1G) to confirm the deletion of the target genes.

### Deletion of *luxS* in *F*. *periodonticum* with *galK* as a counterselection marker

The plasmid pZP06Δ*galK* was introduced into *F. periodonticum* ATCC 33693 competent cells via electroporation. The same procedure was followed for *galK* deletion to obtain plasmid-integrated strains. Once integration was confirmed, one pZP06Δ*galK*-integrated strain was cultured overnight in TSPC broth without antibiotics. The following day, the culture was diluted 1,000-fold and 10,000-fold into fresh TSPC medium, and 100 µL of each dilution was spread onto TSPC agar plates containing 0.25% 2-DG. This medium allowed the selection of cells where the integrated plasmid had been removed through a second recombination event. After three days of growth in an anaerobic chamber, approximately 20 colonies were selected. To assess their antibiotic sensitivity, these colonies were individually re-streaked onto both regular TSPC plates and TSPC plates supplemented with 5 µg/mL thiamphenicol. All colonies exhibited sensitivity to thiamphenicol, indicating the loss of the integrated plasmids. Ten of these colonies were further analyzed by PCR using the primer pair P3/P4 to confirm the deletion of the *luxS* gene.

### Deletion of *radD* gene in fusobacterial strains with *sacB* as a counterselection marker

The procedure for markerless deletion of the *radD* gene was similar to that used for *galK* and *luxS*, including plasmid integration and selection. Once plasmid-integrated strains were confirmed, they were subjected to counterselection on TSPC agar plates with sucrose (5% for ATCC 23726 and ATCC 10953; 10% for CTI-2 and 21-1A) to identify strains that had lost the plasmid through a second recombination event. Thiamphenicol-sensitive colonies were then PCR-verified for *radD* deletion with primers P7/P8.

### AI-2 assay

The AI-2 bioluminescence assay was conducted following established protocols (54). In brief, bacterial cultures in the stationary phase were centrifuged at 12,000 g for 5 minutes to pellet the cells. The resulting supernatant was filtered through a 0.22 μm filter (Millipore, Bedford, MA, USA) to obtain cell-free supernatant (CS) samples. The reporter strain *Vibrio harveyi* BB170 was diluted 1:5000 in fresh autoinducer bioassay medium (consisting of 0.3 M NaCl, 0.05 M MgSO₄, 0.2% Casamino Acids, 10 μM KH₂PO₄, 1 μM L-arginine, 20% glycerol, 0.01 μg/mL riboflavin, and 1 μg/mL thiamine) and cultured at 30°C for 3–4 hours. After incubation, 180 μL of the *V. harveyi* culture was mixed with 20 μL of the CS sample and incubated at 30°C for 4 hours. After incubation, 100 μL aliquots were transferred to a black flat-bottomed 96-well microplate (Greiner Bio-one: #655096) for bioluminescence measurement using a GloMax® Navigator Microplate Luminometer. Cell-free supernatant from *Escherichia coli* BL-21 was the positive control, while its *luxS* mutant was the negative control. The *V. harveyi* bioassay was performed in triplicate for each sample within a single experiment, and the experiments were repeated three times.

### Bacterial co-aggregation

Using a previously described method, co-aggregation assays were conducted using *F. nucleatum* wild-type strains and Δ*radD*, combined with *A. oris* MG-1. In brief, stationary-phase cultures of bacterial strains were grown in TSPC with/ without inducers or heart infusion broth for MG-1. Post centrifugation, cells were washed and resuspended in coaggregation buffer (200 mM Tris-HCl, pH 7.4, 150 mM NaCl, 0.1 mM CaCl_2_), ensuring an equal cell density of approximately 2 x 10^9^ ml^-1^ based upon OD_600_ values. For the assay, 0.20 ml aliquots of *Actinomyces* and various fusobacterial cell suspensions were mixed in a 12-well plate, briefly shaken on a rotator, and then imaged.

### Western blotting analysis

Previously, we generated an antibody against RadD using a recombinant protein corresponding to amino acids 41-360 in the N-terminal region of RadD from ATCC 23726 (55). However, this antibody exhibited nonspecific reactivity with unknown proteins (55). To improve specificity, we developed a new RadD antibody using a recombinant protein containing amino acids 44-200 of RadD from ATCC 23726, a region highly conserved among RadD homologs (Fig. S1). The RadD fragment was amplified using primers EX-radD-E/F and cloned into the pMCSG53 vector. This construct was transformed into *E. coli* BL21(DE3), and the His6-tagged RadD protein was purified through affinity chromatography. The purified protein was then used to produce antibodies via Cocalico Biologicals, Inc. For Western blotting, 1 mL overnight cultures of various *Fusobacterium* strains and their respective *radD* mutants were collected, washed twice with water, and resuspended in SDS-PAGE loading buffer. After boiling the samples at 70°C for 10 minutes, they were subjected to 4%-20% Tris-glycine gradient SDS-PAGE. Immunoblotting was performed using the newly produced rabbit anti-RadD antibody and an anti-HsIJ antibody (used as a reference protein) at a 1:1000 dilution.

## AUTHOR CONTRIBUTIONS

P.Z. and C.W. conceived and designed all experiments. P.Z., B.G., and C.W. performed all experiments. P.Z. and C. W. analyzed data. C.W. and P.Z. wrote the manuscript with contribution and approval from all authors.

## DATA AVAILABILITY STATEMENT

Materials are available upon reasonable request with a material transfer agreement with UTHealth for bacterial strains or plasmids. Plasmids information can be accessed on Benchling links at https://benchling.com/s/seq-oKUe1TAUIUjXcZGa47YQ?m=slm-7eQ5N6wPazrDAIFAALlK (pCWU50);

https://benchling.com/s/seq-fd2pWoLq97uSAuCO3BHH?m=slm-DFavm5OORUNc1SwV5b0Y (pZP05);

https://benchling.com/s/seq-b6iEWGFNntRRRXBpLFb8?m=slm-xDSqzrc8PeBJWOYTIhHw (pZP06);

https://benchling.com/s/seq-Ocb2mMpIfBnfdk1J8keP?m=slm-esofSb8BrvcT605TPIzs (pZP06B);

https://benchling.com/s/seq-E645TWtGGB3eGGmoxV06?m=slm-Vbl2a62xQEczyAUCHKd9 (pZP06C).

## ACKNOWLEDGMENTS

This work was supported by NIDCR grants DE030895 and DE034542 awarded to C.W. We are grateful to Dr. Wendy Garrett (Harvard T.H. Chan School of Public Health) and Dr. Daniel Slade (Virginia Tech) for providing the CTI-2 and 21_1A strains, both of which were sourced from CRC samples. We also thank Dr. Xuesong He (The Forsyth Institute) for supplying *Vibrio harveyi* BB152 and BB170 and Dr. Xiangan Han (Shanghai Veterinary Research Institute, Chinese Academy of Agricultural Sciences) for providing the *E. coli* BL-21 (DE3) Δ*luxS* strain.

